# CrusTome: A transcriptome database resource for large-scale analyses across Crustacea

**DOI:** 10.1101/2022.11.03.515067

**Authors:** Jorge L. Pérez-Moreno, Mihika T. Kozma, Danielle M. DeLeo, Heather D. Bracken-Grissom, David S. Durica, Donald L. Mykles

**Author notes:** Corresponding author: Jorge L. Pérez-Moreno.

## Abstract

Transcriptomes from non-traditional model organisms often harbor a wealth of unexplored data. Examining these datasets can lead to clarity and novel insights in traditional systems, as well as to discoveries across a multitude of fields. Despite significant advances in DNA sequencing technologies and in their adoption, access to genomic and transcriptomic resources for non-traditional model organisms remains limited. Crustaceans, for example, being amongst the most numerous, diverse, and widely distributed taxa on the planet, often serve as excellent systems to address ecological, evolutionary, and organismal questions. While they are ubiquitously present across environments, and of economic and food security importance, they remain severely underrepresented in publicly available sequence databases. Here, we present CrusTome, a multi-species, multi-tissue, transcriptome database of 201 assembled mRNA transcriptomes (189 crustaceans, 30 of which were previously unpublished, and 12 ecdysozoan outgroups) as an evolving, and publicly available resource. This database is suitable for evolutionary, ecological, and functional studies that employ genomic/transcriptomic techniques and datasets. CrusTome is presented in BLAST and DIAMOND formats, providing robust datasets for sequence similarity searches, orthology assignments, phylogenetic inference, etc., and thus allowing for straight-forward incorporation into existing custom pipelines for high-throughput analyses. In addition, to illustrate the use and potential of CrusTome, we conducted phylogenetic analyses elucidating the identity and evolution of the Cryptochrome Photolyase Family of proteins across crustaceans.

## Introduction

A distinct paucity of readily available genomic and transcriptomic resources persists for non-model organisms, despite recent advances in sequencing technologies and in the adoption of bioinformatics across diverse fields (Mykles *et al*. 2016; Burnett *et al*. 2020). These non-model organisms often harbor a wealth of potentially useful genomic and transcriptomic data, which can lead to discoveries and unforeseen advances in a diverse array of seemingly unrelated areas (GIGA Community of Scientists 2014; Tagu *et al*. 2014). Crustaceans are amongst the most numerous and diverse taxa on the planet (Martin and Davis 2006; Ahyong *et al*. 2011; Schram 2013). Thanks to their ubiquitous presence across an extreme diversity of biomes (Pérez-Moreno *et al*. 2016; Bracken-Grissom and Wolfe 2020), they are particularly well suited to address questions of ecological, evolutionary, and organismal interest (Stillman *et al*. 2008; Pérez-Moreno *et al*. 2018; Wolfe *et al*. 2021). In addition to their critical environmental and scientific relevance, crustaceans are of major significance for social, economic, and food security implications (Timm *et al*. 2019; Boyd *et al*. 2022). Nevertheless, similar to other non-model invertebrates, they are severely underrepresented in publicly accessible (and readily available) databases such as those maintained by the National Center for Biotechnology Information (NCBI) (GIGA Community of Scientists 2014; Havird and Santos 2016; Hyde *et al*. 2020). Furthermore, obtaining data from raw data databases (such as the NCBI’s Sequence Read Archive) and transforming them into a useable format represents a time-consuming and computationally expensive process, ultimately hindering accessibility by non-specialists and researchers with limited computational, temporal, or financial resources.

Here we present CrusTome (from *crustacean* and Greek *tomos*, book or volume), a multi-species, multi-tissue, transcriptome database of 201 assembled mRNA transcriptomes (at the present time 189 crustaceans, 30 of which were previously unpublished, and 12 additional outgroups from among Ecdysozoa) to aid in evolutionary inference and as support for evolutionary, ecological, and functional studies that employ genomic/transcriptomic techniques for sequence similarity searches, orthology assignments, and phylogenetic inference, among other uses. This CrusTome database is initially presented in BLAST and DIAMOND formats for convenience and ease of use, aiming to provide robust datasets for sequence similarity searches, orthology assignments, phylogenetic inference, among other uses. Presenting the database in this format also allows for simple and straight-forward incorporation into existing custom analysis pipelines, making it particularly suitable for scripting and high-throughput analyses (e.g., Pérez-Moreno *et al*. 2018; Drozdova *et al*. 2021). The database will be updated regularly with new data as new species and tissues are sequenced, as well as when advances in bioinformatic pipelines warrant the re-processing or re-assembly of the raw reads. This will be achieved with a high-memory computing node located at the OU Supercomputing Center for Education & Research (OSCER) at the University of Oklahoma (OU). In addition, we present an example in which we employ the CrusTome database to conduct the first large-scale transcriptomic exploration across crustaceans of the Cryptochrome/Photolyase Family (CPF) – a class of light-sensitive flavoproteins involved in DNA repair, circadian rhythm regulation, and magnetoreception that have also shown promising applications as optogenetic tools (Oliveri *et al*. 2014; Mei and Dvornyk 2015; Hernández-Candia and Tucker 2020; Kiontke *et al*. 2020). Links to download the CrusTome database and associated metadata, future updates, and the code to reproduce the example analysis hereby presented, can be found at CrusTome’s GitHub site: https://github.com/invertome/crustome

## Methods

### Data sourcing

For methodological consistency and reliability of downstream analyses, transcriptomes were assembled from raw RNAseq reads, when these were publicly available. Emphasis was placed on raw reads of non-hexapod *Pancrustacea* samples (n=189) covering the phylogenetic breadth available on the NCBI’s SRA database (excepting Hexapoda; Leinonen *et al*. 2011), with a minimum library requirement of 1M reads in depth per sample downloaded (Figure 1A). Additional raw reads of ecdysozoan samples (n=12) were included to serve as outgroups during gene characterization and phylogenetic inference. Sequencing reads of nematodes from the family Monhysteridae were also included as species from this family are frequently found as parasites or in association with numerous crustacean species (Baylis 1915; Chitwood 1935; Westerman *et al*. 2022; Tchesunov and Ivanenko), and were thus deemed useful as a mechanism to filter out potential non-crustacean contaminant sequences during downstream analyses. Full details on the transcriptomes included, Accession Identifiers, and corresponding raw read and transcriptome metadata can be found in Supplementary File 2.

**Figure 1.**
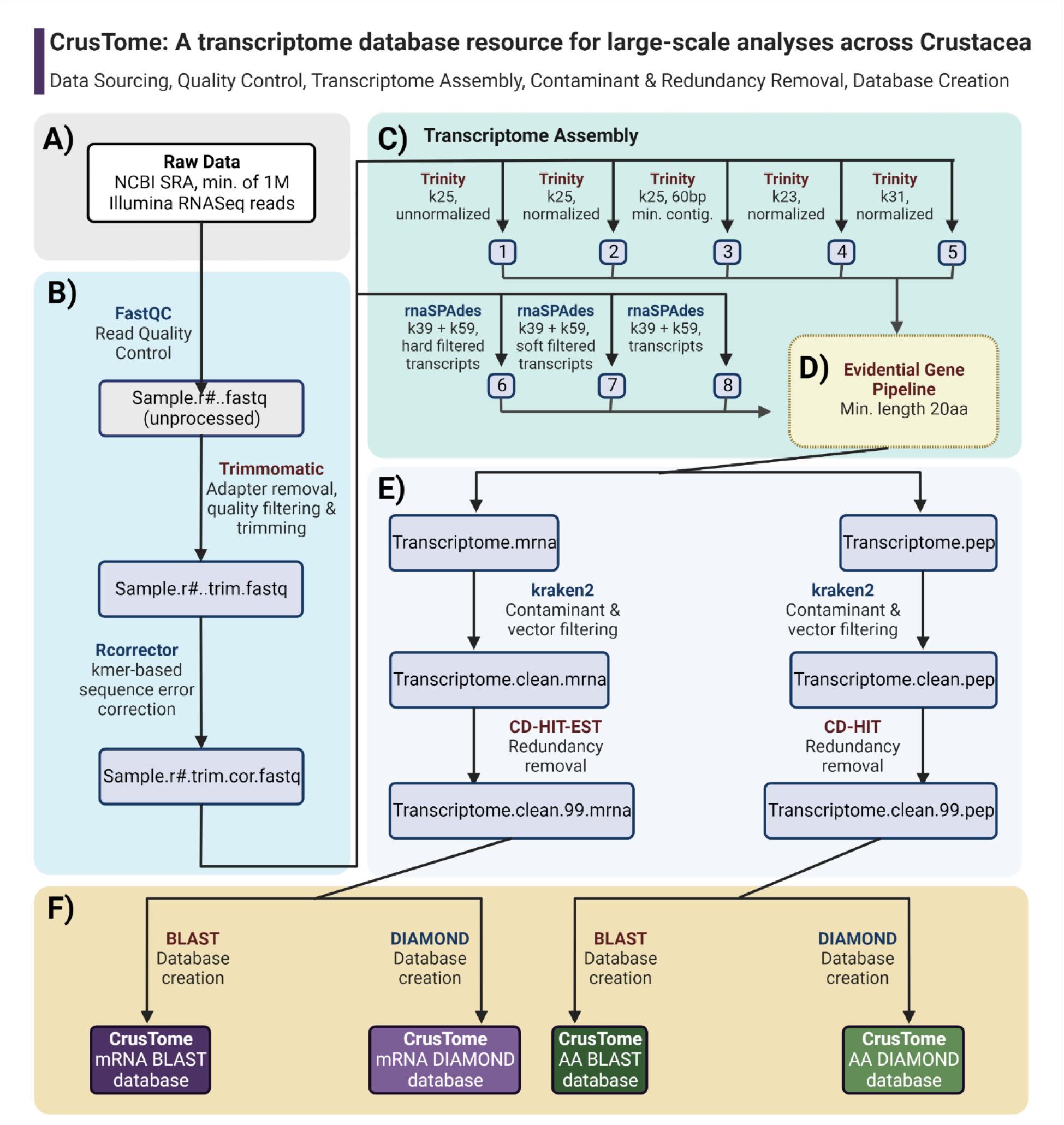
Flow-chart of the CrusTome database creation steps. A. Data sourcing; B. Quality Control and Pre-Processing; C. Transcriptome Assembly with multiple assemblers; D. Obtaining optimal transcriptome via EvidentialGenePipeline; E. Contamination and Redundancy Filtering; F. Creation of BLAST and DIAMOND databases.

### Quality Control

Sequencing reads downloaded from the NCBI’s SRA database were visually inspected using FastQC (Andrews 2010) to determine read filtering, trimming stringency, and thresholds. As samples were of a heterogeneous nature with a range of input qualities, trimming settings were set up conservatively to ensure proper transcriptome assemblies and the reliability of downstream analyses. Automated trimming of the sequencing reads was undertaken using Trimmomatic (Bolger *et al*. 2014) with the following settings: *CROP=“x” HEADCROP=“15” MINLEN=“45” SLIDINGWINDOW=“4:20” LEADING=“15” TRAILING=“15”*, with the *CROP* value “x” adjusted to remove the error-prone final 10 error-prone bases according to each sequencing library’s fragment length (e.g., 90 for 100 bp reads, or 140 for 150 bp reads). Immediately subsequent to trimming, the resulting sequencing read files were piped into Rcorrector (using default settings) for k-mer based random sequencing error correction using a De Bruijn graph algorithm, which is particularly suitable for error-correction of RNAseq reads (Song and Florea 2015; Figure 1B).

### Transcriptome Assembly

The quality filtered, trimmed, and error corrected reads were assembled into *de novo* transcriptomes for each sample using a multi-assembler approach (Figure 1C) that leverages the advantages of different assembly algorithms and parameters to obtain a single optimal, and less fragmented, transcriptome assembly (Nakasugi *et al*. 2014; Gilbert 2019). Transcriptomes were initially assembled via several iterations of the Trinity pipeline (version 2.13.2; Grabherr *et al*. 2011; Haas *et al*. 2013), with differing k-mer, normalization, and minimum contig-length parameters (*k25 + unnormalized reads, k25 + normalized reads, k25 + 60 bp min contig length, k31 + normalized, k23 + normalized*), exclusively retaining transcripts supported by read mappings (Haas *et al*. 2013). Additional assemblies were produced using rnaSPAdes (version 3.15.13; Bankevich *et al*. 2012) and its integrated multi-kmer assembly approach. rnaSPAdes was run with two k-mer settings (k39 + k59) after which assembled contigs were collated into three assemblies resulting from differing quality filtering thresholds (*hard_filtered_transcripts*.*fasta, soft_filtered_transcripts*.*fasta, transcripts*.*fasta*), all of which were included in subsequent merging steps. All of the aforementioned resulting Trinity and rnaSPAdes assemblies (n=8) for each sample were then merged into a single optimal transcriptome via the EvidentialGene pipeline (EVG; Gilbert 2019), with a selected minimum amino acid length of 20 residues (e.g., to allow for the detection and characterization of small neuropeptides; Figure 1D). This multi-assembler and multi-parameter scheme was empirically determined to obtain transcriptome assemblies of higher quality and completeness, by capitalizing on the advantages of each set of assembler/parameters, as determined by summary statistics and BUSCO scores (Simão *et al*. 2015) among other metrics of protein completeness (Gilbert 2019). Final merged transcriptomes obtained were filtered with the EVG pipeline for long non-coding RNA (lncRNA) sequences, which are beyond CrusTome’s current scope, to improve the efficiency of sequence similarity searches and downstream analyses. The filtered mRNA transcriptomes were then translated into amino acid sequences to produce both transcribed and translated versions for convenience and accessibility. Samples from previously assembled, unpublished data were included in their original versions and processed through the EVG pipeline for consistency and suitability of comparisons. It is important to note that these samples continue to be subject of reassembly and will be included in their final multi-assembler versions in upcoming releases of CrusTome. Transcriptome assembly and associated bioinformatic analyses were performed using computational resources provided by the Instructional & Research Computing Center (IRCC) at Florida International University and the Supercomputing Center for Education & Research (OSCER) at the University of Oklahoma.

### Contaminant filtering & redundancy removal

Both mRNA and amino acid transcriptomes for each sample were subsequently filtered for contamination using Kraken 2.1.2 (Wood *et al*. 2019; Wright *et al*. 2022) and a custom database prepared in-house that includes archaea, bacteria, virus, fungi, mouse, human, and sequencing vector sequences (sequences obtained from https://lomanlab.github.io/mockcommunity/mc_databases.html and Bushnell, 2018) with settings optimized for filtering crustacean transcriptomes (“*--confidence 0*.*1*”; Wright *et al*. 2022). It is important to note that the confidence setting employed was empirically determined in an iterative process following Wright *et al*. (2022) to filter out most contaminant sequences with minimal loss of crustacean sequences. However, it is possible that some non-crustacean sequences from symbionts may still be present in the database: further filtering steps according to each specific scenario are encouraged (e.g., via a phylogenetic assessment). A final application of the CD-HIT-EST and CD-HIT clustering algorithm was run on each individual EVG-optimized transcriptome (mRNA and amino acid, respectively) to cluster sequence at a 99% sequence identity similarity (Figure 1E). This allowed for the removal of contigs likely to have arisen from sequencing errors, while minimizing the removal of true isoforms.

### Transcriptome summary statistics & completement assessment

TransRate (1.0.3) and Benchmarking Universal Single-Copy Orthologs (BUSCO) 3.0.2 (Simão *et al*. 2015; Smith-Unna *et al*. 2016) were employed to calculate summary statistics and to assess the completeness of the CrusTome transcriptome assemblies. BUSCO’s software evaluates transcriptome completeness in an evolutionary informed context by assessing the presence and/or fragmentation of universal single copy orthologs (Simão *et al*. 2015). BUSCO analyses were conducted using OrthoDB’s Arthropoda database of orthologous groups (Waterhouse *et al*. 2013) as a reference dataset.

### BLAST & Diamond database creation

The CrusTome mRNA and amino acid databases were created in two formats using default settings, as both BLAST and Diamond databases (Altschul *et al*. 1990; Buchfink *et al*. 2021; Figure 1F) for compatibility with existing annotation and analysis pipelines (e.g., see Das *et al*. 2016; Pérez-Moreno *et al*. 2018; Tang *et al*. 2019; Drozdova *et al*. 2021) and upcoming software. Diamond is an ultra-fast alignment software that achieves considerable sequence similarity search speeds orders of magnitude faster than BLAST, at a minimum sensitivity cost (Buchfink *et al*. 2021), and as such would be appropriate for the high-throughput applications now available with CrusTome.

### “CrusTome” example pipeline

An example analysis was conducted to illustrate the potential of CrusTome for the identification and characterization of peptides of interest. Specifically, we conducted a series of recursive BLAST searches against CrusTome’s amino acid sequence database, followed by an alignment and phylogenetic inference strategy to gain insight into the presence and expression of DNA-photolyases, Cryptochromes, and DASH-like Cryptochromes (Oliveri *et al*. 2014; Mei and Dvornyk 2015; Kiontke *et al*. 2020) across crustaceans.

The phylogenetically informed annotation analyses consisted of an initial BLAST search against CrusTome using reference Cryptochrome and DNA-photolyase sequences previously characterized in insects (Supplementary File 1), specifying a maximum number of hits of 500 to capture as much sequence diversity as possible (Shah *et al*. 2019), but with a relatively stringent e-value of e^-120^ to limit results to relevant peptides. The list of hit IDs resulting from this initial search was then used to extract the corresponding sequences from CrusTome, which were then used as input for a second BLAST search against the database to capture additional sequence diversity. Sequences identified as hits from this second BLAST iteration were once again extracted from CrusTome. These putative peptides identified by BLAST were subsequently concatenated with the insect references originally employed as search queries, which were then aligned with the multiple sequence aligner MAFFT (v.7.490; Yamada *et al*. 2016). The MAFFT software was then invoked to align putative Cryptochrome and Photolyase sequences obtained from the CrusTome database, along with the original insect reference sequences used as BLAST queries. MAFFT alignment parameters were specifically chosen to prioritize accuracy over speed and to allow for large unalignable regions that can be pervasive in certain protein families (“ *-- dash --ep 0 --genafpair --maxiterate 1000*”, as in Pérez-Moreno *et al*. 2018). Additionally, the -- *dash* parameter enables MAFFT to query a Database of Aligned Structural Homologs (DASH), providing structural information with which to refine the alignment process (Rozewicki *et al*. 2019). Following alignment completion, the resulting alignment was then trimmed using ClipKit (Steenwyk *et al*. 2020), which identifies and retains phylogenetically informative sites for a more accurate and robust phylogenetic inference. Phylogenetic reconstruction was undertaken with IQ-tree (Nguyen *et al*. 2015) with a Le-Gascuel (LG) general amino acid replacement matrix under a FreeRate model with 10 rate categories (LG+R10; Yang 1995; Müller and Vingron 2000; Le and Gascuel 2008; Soubrier *et al*. 2012) as suggested for our data by ModelFinder (Kalyaanamoorthy *et al*. 2017). The phylogenetic tree resulting from this initial reconstruction was then piped, in conjunction with the alignment, to TreeShrink for outlier/paralog detection and removal at an α value of 0.05 (Mai and Mirarab 2018). The resulting pruned alignment was then used for a second and final phylogenetic reconstruction with IQ-tree (Nguyen *et al*. 2015) for further characterization and annotation of the putative peptides. The second IQ-tree phylogenetic reconstruction was run using the same model parameters previously reported (LG+R10; Yang 1995; Müller and Vingron 2000; Le and Gascuel 2008; Soubrier *et al*. 2012). Branch support of this final phylogeny was assessed in bipartite by Ultra-fast bootstrap approximation (UFBoot; 10,000 replicates) and an approximate Bayes test (Guindon *et al*. 2010; Anisimova *et al*. 2011; Minh *et al*. 2013). Finally, the resulting phylogenies were used to putatively classify the obtained peptide sequences as members of Cryptochrome 1, Cryptochrome 2, DASH-like Cryptochromes, 6-4 Photolyases, or CPD photolyases as per previous studies in other organisms (Oliveri *et al*. 2014; Mei and Dvornyk 2015; Kiontke *et al*. 2020). Protein sequences were collated for each of these major clades, and each of the sequence groups were then aligned following the previously mentioned strategy. The gene-specific alignments were then used to generate Hidden Markov Model profiles with HMMER (Finn *et al*. 2011; Eddy 2011). These profiles are made available with CrusTome as a community resource. Example code for this phylogenetic analysis is included as Supplementary File 2.

## Results & Discussion

The underrepresentation of non-hexapod pancrustaceans in publicly available databases could be attributed to challenges of a technical nature rather than to a lack of effort or adoption of genomic methodologies by researchers. This disparity is exemplified by the rapid increase of raw sequencing reads in the NCBI’s Sequence Read Archive (SRA) (Havird and Santos 2016; Qin *et al*. 2017; Hyde *et al*. 2020), in contrast to the Transcriptome Shotgun Assembly (TSA) database. Since the SRA database is comprised of raw sequencing reads, accessing information stored therein requires a specific expertise and set of skills, oftentimes with steep learning-curves. Given that crustacean ‘omics data are now being produced at a far greater rate than can be meaningfully accessed, analyzed and interpreted by many researchers, CrusTome delivers a solution that is simple to implement to enable large scale transcriptomic analyses across non-hexapod Pancrustacea. The multiple k-mer assembly strategy and subsequent merging through the EVG pipeline, as employed in the preparation of CrusTome, offers noticeable advantages for *de novo* transcriptome assembly from organisms without a reference genome (Gilbert 2019; Summary Statistics & BUSCO Assessment, Supplementary File 3). Additionally, special emphasis was placed on the processing of the publicly available data by having re-assembled and processed each included transcriptome with the aforementioned pipeline rather than including assemblies produced by disparate methodologies. This processing ensures increased contiguity and decreased fragmentation of the resulting assemblies, as well as their standardization for accurate and consistent downstream analyses.

Several approaches have been previously taken to address this knowledge gap representing crustaceans and other non-traditional model organisms (Qin *et al*. 2017; Nong *et al*. 2020; Hyde *et al*. 2020). For example, CrusTF is a web-based database resource containing sequence data, with an emphasis on transcription factors, mined from multiple transcriptomic sources (Qin *et al*. 2017). Having sourced data from over 170 transcriptomes, CrusTF is the most taxonomically diverse curated database available to date. However, despite the taxonomic breadth covered across Crustacea, ease of access, and web-based tools and operability, its present scope is limited, as it pertains exclusively to transcription factors. CrustyBase, a recently published interactive database of crustacean transcriptomes, also employs a web-based approach with a Graphical User Interface (GUI) that excels in terms of accessibility, navigation, and operability (Hyde et al., 2020). It also leverages the advantages of being able to process gene expression data that can be linked directly to each submitted transcriptome, an integrated BLAST interface, and intuitive visualization features. Nevertheless, although highly curated, it is dependent upon direct submissions from the community, and suffers from underrepresentation of numerous crustacean taxa. Furthermore, similar to CrusTF, CrustyBase’s main target audience consists of those comfortable with conducting analyses exclusively through GUIs. While that presentation is certainly of advantage for data accessibility to a specific sector that may be unfamiliar with coding, it hinders utilization by those who wish to incorporate their datasets into existing command-line based bioinformatic pipelines for large-scale and/or high-throughput analyses such as those now made possible by CrusTome.

CrusTome’s current version consists of a multi-species and multi-tissue transcriptome database from 189 non-hexapod pancrustacean species, including 30 previously unpublished transcriptomes, and 12 additional ecdysozoan species (see Supplementary File 3 for additional details). This initial version of CrusTome includes a sample of resources currently available on the NCBI’s public repositories and therefore is subject to similar representation biases (Figure 2A). Consequently, CrusTome should be considered an evolving database resource, as it will continue to be updated to bridge these gaps whenever relevant data become available. The present version presents an uneven distribution of samples across Pancrustacean taxa biased towards the class Malacostraca, which comprises 174 out of 201 transcriptomes (Figure 2B). Despite this apparent overrepresentation, CrusTome includes representative of rare and obscure taxa that present intriguing opportunities for phylogenetics, systematics, and evolution, such as remipedes and bathynellaceans (Pérez-Moreno *et al*. 2016) and multiple deep-water malacostracans (DeLeo and Bracken-Grissom 2020; Drozdova *et al*. 2021). Samples of pancrustacean taxa Ostracoda, Mystacocarida, Branchiura, and Cephalocarida are currently in the processing queue for upcoming iterations, improving CrusTome’s phylogenetic breadth. In addition, other members of Pancrustacea (namely hexapods), as well as a select number of Chelicerata and Tardigrada transcriptomes have been included to aid in comparative analysis and serve as outgroups to root phylogenies, along with nematodes from the family Monhysteridae to filter potential contamination from symbiotic organisms and/or parasites (Baylis 1915; Westerman *et al*. 2022). In addition to phylogenetic diversity, CrusTome also provides a wide array of sample types, from single tissues to whole organisms, aiming to encompass transcript diversity both across and within species.

**Figure 2.**
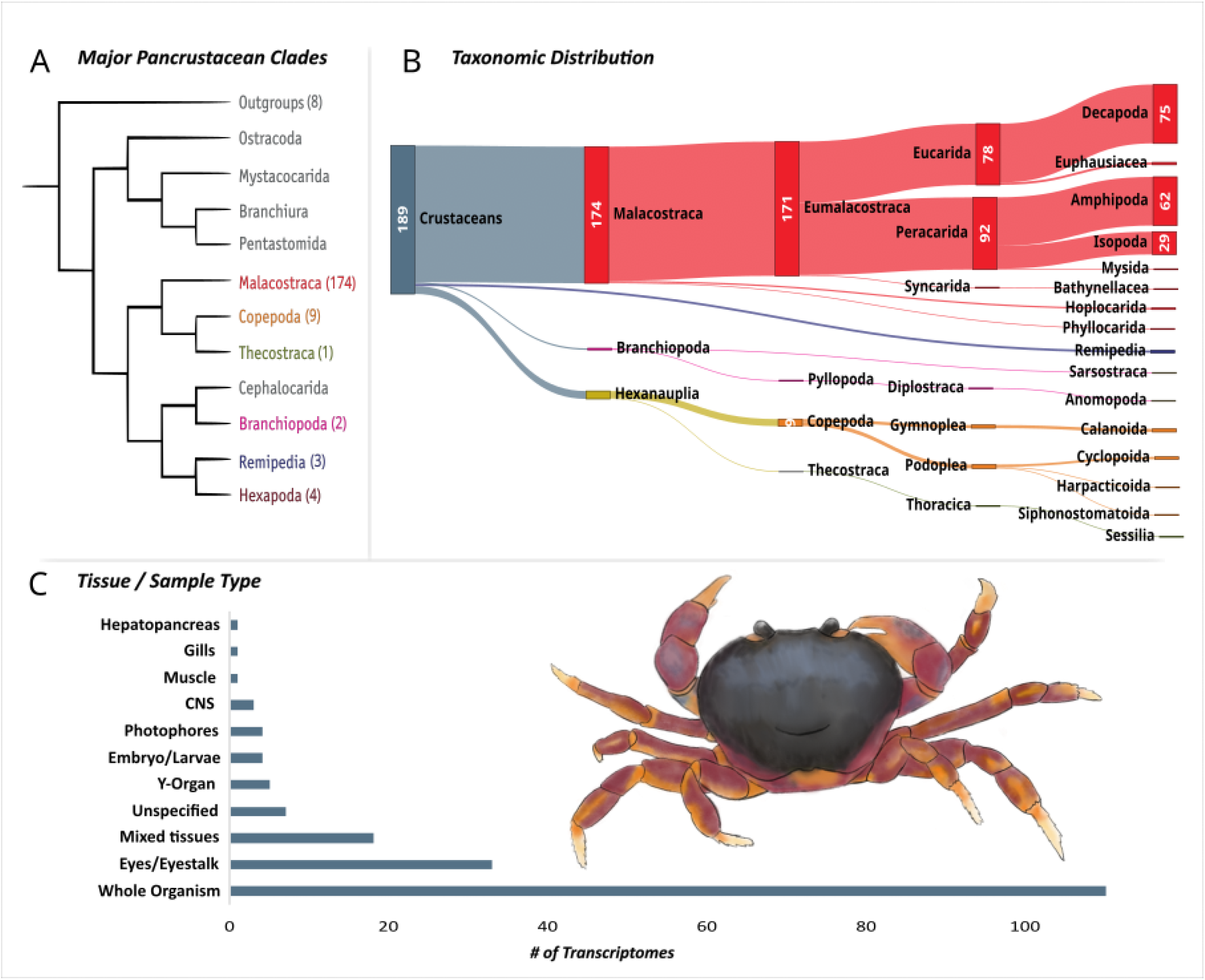
A) Taxon coverage across major Pancrustacean clades in the current version of the CrusTome database. Phylogeny adapted from Oakley *et al*. 2013; Bracken-Grissom and Wolfe 2020). B) Sankey diagram depicting the taxonomic distribution of transcriptomes included in the present version of the database. C) Tissue type distribution of CrusTome transcriptomes across non-hexapod pancrustaceans. Illustration of *Gecarcinus lateralis* by An-Ping Yu.

### Example Analysis: Large-scale exploration of Cryptochromes and DNA-Photolyases across Crustacea

To illustrate the functionality of the CrusTome database, an analysis was conducted to characterize putative Cryptochrome/Photolyase Family proteins expressed across multiple tissues from species spanning the pancrustacean phylogenetic tree. Cryptochromes and photolyases are UV-A/blue-light sensitive proteins that can be found across the entire tree of life and share a common general structure of a conserved photosensory domain bound to two chromophore cofactors (Sancar 2003; Chaves *et al*. 2011; Oliveri *et al*. 2014; Mei and Dvornyk 2015). However, important functional differences exist between the two types of CPF proteins. Photolyases are light-dependent DNA-repair enzymes that can be classified based on the type of damage they repair: (1) the cyclobutene pyrimidine dimer (CPD) photolyases and (2) the 6-4 pyrymidine-pyrimidone photoproducts (6-4) photolyase (Sancar 2003, 2008; Hitomi *et al*. 2009; Oliveri *et al*. 2014). Despite their structural similarity with photolyases, cryptochromes are not involved in DNA repair activity, and instead participate in a wide variety of functions, such as light perception, transcriptional regulation, and magnetoreception (Chaves *et al*. 2011; Liu *et al*. 2011; Oliveri *et al*. 2014; Bazalova *et al*. 2016). Although CPF proteins are known to be present in all types of organisms (prokaryotic and eukaryotic; Reitzel *et al*. 2010; Rivera *et al*. 2012; Zantke *et al*. 2013), including crustaceans (i.e., the copepod *Eurydice pulchra* and the Antarctic Krill, *Euphausia superba*; Teschke *et al*. 2011; Zhang *et al*. 2013), but little is known about their distribution and function across Pancrustacea.

Sequence similarity searches with BLAST (Altschul *et al*. 1990), using reference CPF sequences from the NCBI GenBank (sequences and Accession IDs can be found in Supplementary File 1), recovered putative CPF proteins from CrusTome’s amino acid sequence database. A total of 382 unique sequences were obtained from the 201 transcriptomes included in the database. These sequences were subsequently aligned and trimmed, then used for phylogenetic reconstruction.

The phylogram represented five major CPF clades, which corresponded CPD Photolyases, Cryptochrome 1, Cryptochrome 2, CRY-DASH, and (6-4) Photolyases (Figure 3), whose phylogeny was in overall agreement with previous work (Lin and Todo 2005; Lucas-Lledó and Lynch 2009; Mei and Dvornyk 2015). All of these groups formed monophyletic clades, with the exception of the (6-4) Photolyases, which interestingly fell in their entirety as a clade within Cry2 sequences (Supplementary Materials S4). This confirmed a CPF phylogenetic analysis that found (6-4) Photolyase and Cryptochrome sequences cluster together, in contrast with other homologs (Mei and Dvornyk 2015). Differences in the taxonomic distribution of the five major clades are immediately evident, particularly between the two Cryptochromes. Cryptochrome 1 had a more limited distribution, being found only among amphipods, branchiopods, copepods, decapods, euphausiids, and thecostracans (Figure 4), while Cryptochrome 2 was additionally found in isopods, stomatopods, and mysids (Figure 5). However, care should be taken before making evolutionary or functional inferences, as this difference in distribution may reflect the tissue types included in the database. Nevertheless, the analysis shows the ease of application and potential for novel insights found in large-scale transcriptome analyses through the CrusTome database.

**Figure 3.**
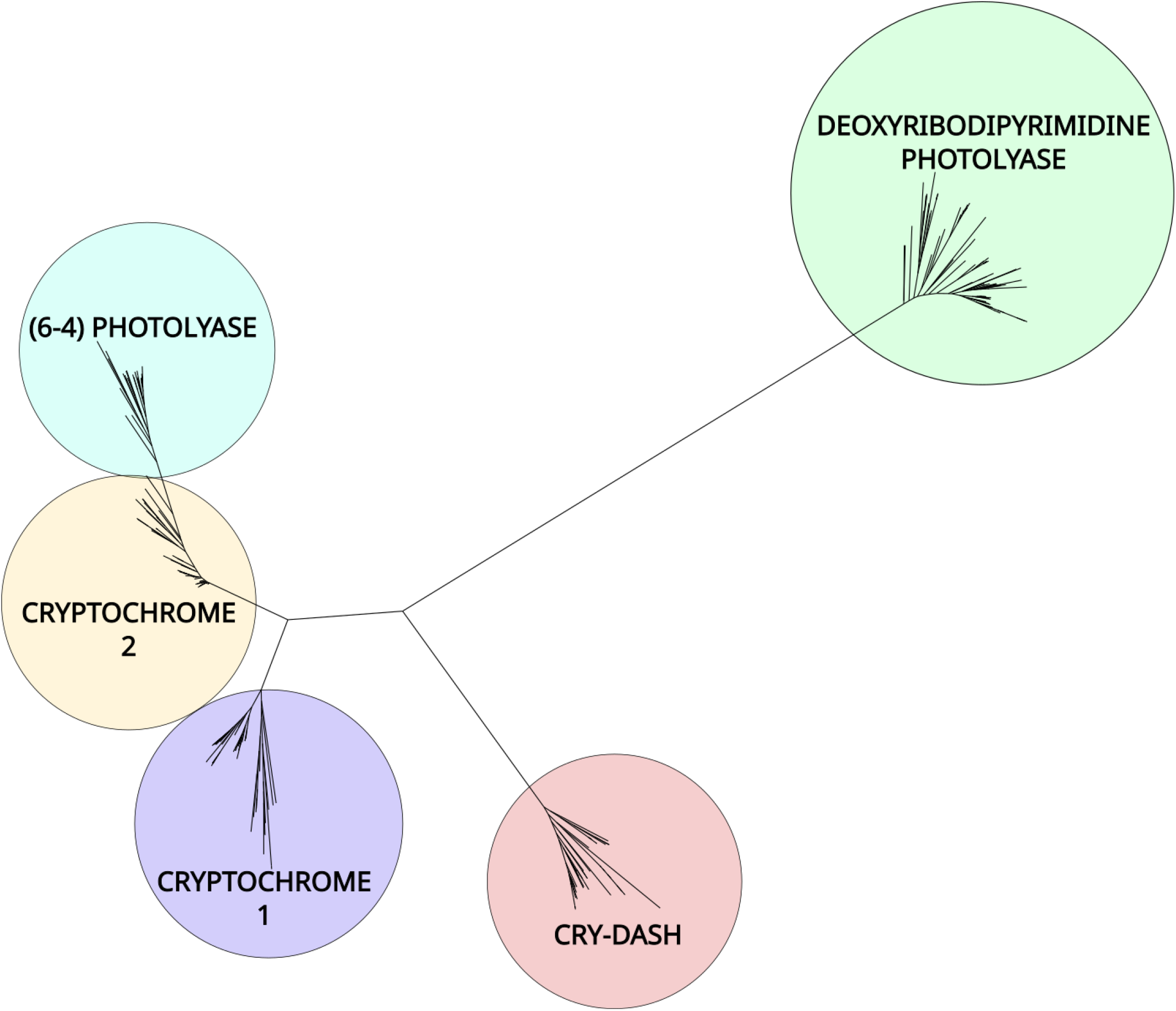
Unrooted phylogenetic tree (LG+R10) of Cryptochrome 1, Cryptochrome 2, Cryptochrome-DASH, and Photolyases in crustacean transcriptomes within the CrusTome database.

**Figure 4.**
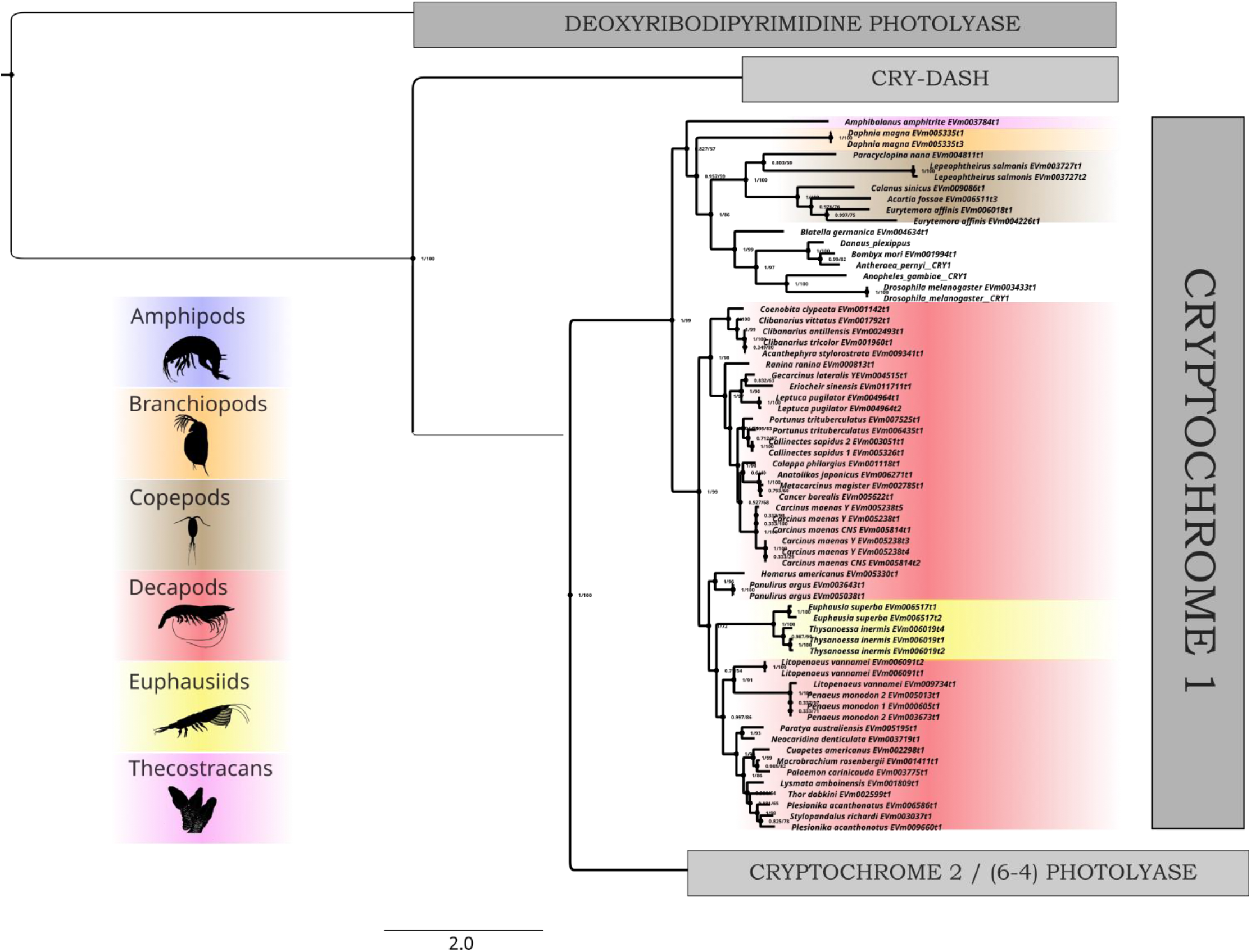
Rooted phylogenetic tree (LG+R10) of Cryptochrome 1 found across transcriptomes of multiple crustacean species and tissues within the CrusTome database. Representative taxa images from PhyloPic.org (Amphipod & decapod by Christoph Schomburg, copepod & thecostracan by Joanna Wolfe, euphausiid by Steven Haddock).

**Figure 5.**
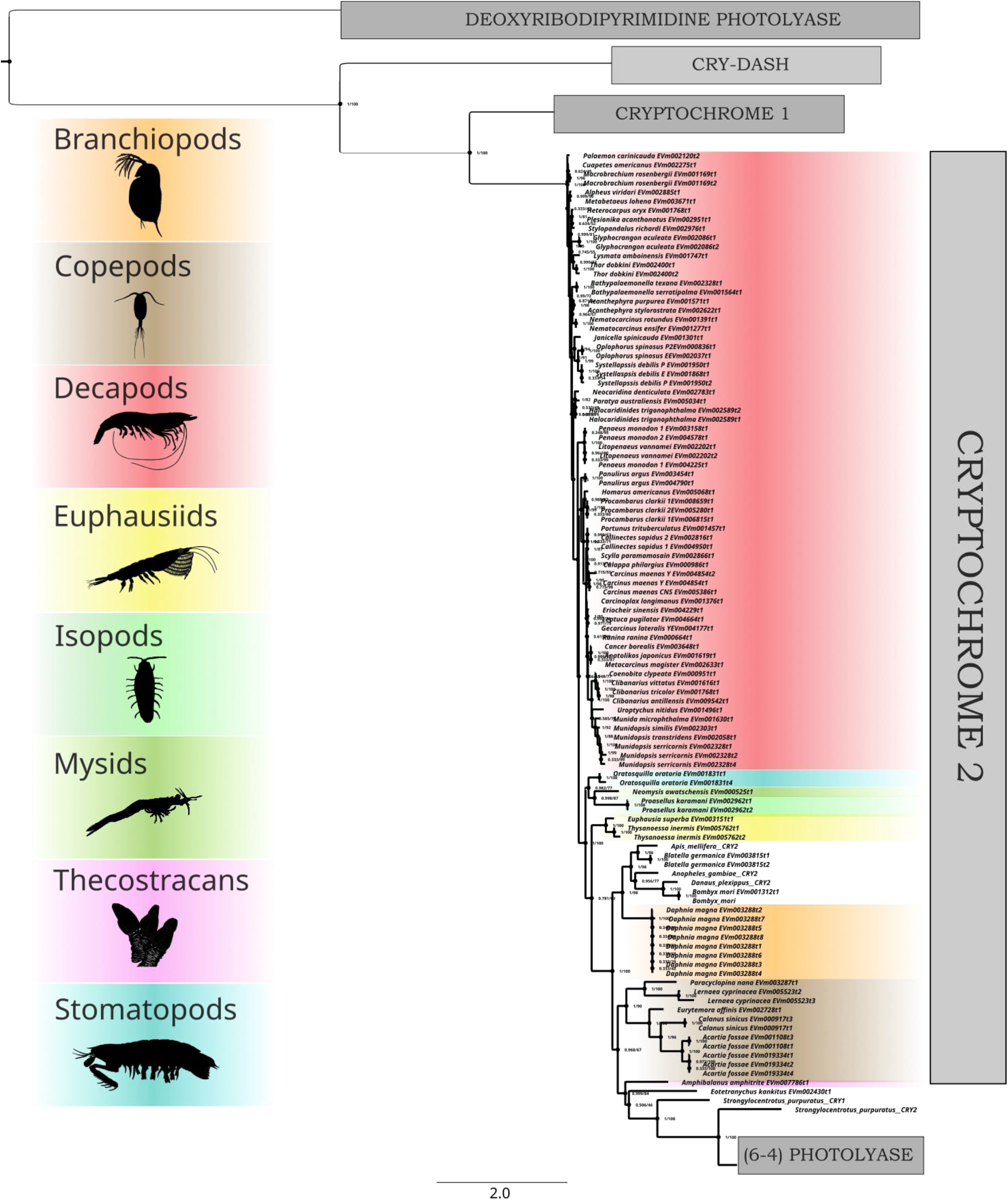
Rooted phylogenetic tree (LG+R10) of Cryptochrome 2 found across transcriptomes of multiple crustacean species and tissues within the CrusTome database. Representative taxa images from PhyloPic.org (Amphipod & decapod by Christoph Schomburg, copepod & thecostracan by Joanna Wolfe, euphausiid by Steven Haddock, isopod by Kanchi Nanjo, mysid by Lafage https://creativecommons.org/licenses/by/3.0/, stomatopod by T. Michael Keesey).

### Future directions and applications

Biased taxon representation in public data repositories is a pressing issue for numerous fields within pancrustacean biology. CrusTome addresses this lack of taxonomic diversity by including fully assembled and pre-processed transcriptomes of underrepresented taxa. It is important to note that the taxonomic distribution of CrusTome’s transcriptomes is ultimately dependent on sequence data that is publicly available and thus may be subject to biases. It is for that reason that CrusTome was envisioned as a community resource that will grow and evolve as data are produced and incorporated to continuously address taxonomic and tissue representation gaps.

The authors also look forward to potential future collaborations with the developers of existing database solutions for crustaceans (i.e., CrusTF, CrustyBase, CAT). Since the scope of the aforementioned projects differs from CrusTome’s, it is important to note that future work integrating these different databases, and leveraging the advantages of each, would be of great benefit to researchers using pancrustaceans as model systems.

## Conclusion

CrusTome provides robust crustacean transcriptome databases in easily accessible formats, using consistent methods for increased reliability and comparability of results. A major goal is to provide a mechanism to improve the current paucity of accessible genomic and transcriptomic data for non-model crustaceans. This accessibility and ease of incorporation into existing pipelines enable analyses at larger scales. Moreover, CrusTome can be used to address long-standing questions in crustacean biology, such as molting and growth (Mykles and Chang 2020; Mykles 2021), sensory biology (e.g., vision, chemoreception; Pérez-Moreno *et al*. 2018; Kozma *et al*. 2020), convergent evolution (e.g. carcinization; Wolfe *et al*. 2021; Yang *et al*. 2021), or adaptation to extreme or changing environments (e.g., caves, deep-sea, polar waters; Pérez-Moreno *et al*. 2016; DeLeo and Bracken-Grissom 2020; Andersen *et al*. 2022). It is our hope that the CrusTome database facilitates access to the rapidly growing number of genomes and transcriptomes being sequenced, particularly to those of non-traditional model organisms. As the transition into a post-transcriptomic era takes place, pancrustacean research must take full advantage of the large amounts of data produced by current and emerging technologies. A major aim of CrusTome is to bridge gaps of knowledge among pancrustaceans by including underrepresented taxa. Accessibility to the large amounts of raw data being deposited in public repositories enables scalable and integrative multi-omic analyses that could ultimately lead to novel biological, ecological, and evolutionary insights across the tree of life (Mykles *et al*., 2010).

## Data Availability Statement

The CrusTome databases have been deposited in the Cyverse repository for public access under DOI: XXXX

Supplementary File 3 (S3) contains a spreadsheet with metadata regarding the raw data employed in building the CrusTome database. Among these data, Accession IDs and sample identifiers are included.

Additional links to the CrusTome database’s associated metadata, previous versions, and example analyses (code, alignments, tree files, etc.) are available at the CrusTome GitHub site: https://github.com/invertome/crustome

## Acknowledgements

The authors would like to acknowledge the Instructional & Research Computing Center (IRCC) at Florida International University and the Supercomputing Center for Education & Research (OSCER) at the University of Oklahoma for providing High Performance Computing resources. The authors are also grateful to Colorado State University’s Crab Lab members for their assistance in testing and optimizing the CrusTome database and associated pipeline, and to An-Ping Yu for providing the crustacean illustrations presented here. This is contribution # XXX from the Coastlines and Oceans Division in the Institute of Environment at Florida International University.

## Supplementary Materials

**S1. *Reference invertebrate cryptochrome and photolyase sequences. FASTA file, S1.cryptochromes.fasta***

**S2. *Example code to conduct a phylogenetic analysis of pancrustacean cryptochromes and photolyases using the CrusTome database. Shell script, S2.cryptochromes.sh***

**S3. *Metadata table. Attached as an Excel file, S3.SuppTable.xlsx***

**S4. *Additional phylograms of the CPF proteins. Attached as a Word document, S4.SuppFigs.docx***

**S5. *CPF protein sequences found within CrusTome. FASTA file, S5.CPF.fasta***

## Literature Cited

Ahyong, S., J. Lowry, M. Alonso, R. Bamber, G. Boxshall et al., 2011 Subphylum Crustacea Brünnich, 1772. In : Zhang, Z.-Q. (Ed.) Animal biodiversity: An outline of higher-level classification and survey of taxonomic richness.

Altschul, S. F., W. Gish, W. Miller, E. W. Myers, and D. J. Lipman, 1990 Basic local alignment search tool. Journal of Molecular Biology 215: 403–410.

Andersen, Ø., H. Johnsen, A. C. Wittmann, L. Harms, T. Thesslund et al., 2022 De novo transcriptome assemblies of red king crab (Paralithodes camtschaticus) and snow crab (Chionoecetes opilio) molting gland and eyestalk ganglia - Temperature effects on expression of molting and growth regulatory genes in adult red king crab. Comparative Biochemistry and Physiology Part B: Biochemistry and Molecular Biology 257: 110678.

Andrews, S., 2010 FastQC A Quality Control tool for High Throughput Sequence Data. http://www.bioinformatics.babraham.ac.uk/projects/fastqc/.

Anisimova, M., M. Gil, J.-F. Dufayard, C. Dessimoz, and O. Gascuel, 2011 Survey of Branch Support Methods Demonstrates Accuracy, Power, and Robustness of Fast Likelihood-based Approximation Schemes. Systematic Biology 60: 685–699.

Bankevich, A., S. Nurk, D. Antipov, A. A. Gurevich, M. Dvorkin et al., 2012 SPAdes: A New Genome Assembly Algorithm and Its Applications to Single-Cell Sequencing. J Comput Biol 19: 455–477.

Baylis, H. A., 1915 LI.—Two new species of Monhystera (Nematodes) inhabiting the gill-chambers of land-crabs. Annals and Magazine of Natural History 16: 414–421.

Bazalova, O., M. Kvicalova, T. Valkova, P. Slaby, P. Bartos et al., 2016 Cryptochrome 2 mediates directional magnetoreception in cockroaches. Proceedings of the National Academy of Sciences of the United States of America 113: 201518622–201518622.

Bolger, A. M., M. Lohse, and B. Usadel, 2014 Trimmomatic: a flexible trimmer for Illumina sequence data. Bioinformatics 30: 2114–2120.

Boyd, C. E., A. A. McNevin, and R. P. Davis, 2022 The contribution of fisheries and aquaculture to the global protein supply. Food Sec.

Bracken-Grissom, H., and J. M. Wolfe, 2020 PHYLOGENY WITH EMPHASIS ON CRUSTACEA. Evolution and Biogeography: Volume 8 80.

Buchfink, B., K. Reuter, and H.-G. Drost, 2021 Sensitive protein alignments at tree-of-life scale using DIAMOND. Nat Methods 18: 366–368.

Burnett, K. G., D. S. Durica, D. L. Mykles, J. H. Stillman, and C. Schmidt, 2020 Recommendations for Advancing Genome to Phenome Research in Non-Model Organisms. Integrative and Comparative Biology 60: 397–401.

Bushnell, Brian, 2018 Masked version of hG 19.

Chaves, I., R. Pokorny, M. Byrdin, N. Hoang, T. Ritz et al., 2011 The cryptochromes: blue light photoreceptors in plants and animals. Annu Rev Plant Biol 62: 335–364.

Chitwood, B. G., 1935 Nematodes parasitic in, and associated with, Crustacea, and descriptions of some new species and a new variety. Proceedings of the Helminthological Society of Washington 2: 93–96.

Das, S., S. Shyamal, and D. S. Durica, 2016 Analysis of Annotation and Differential Expression Methods used in RNA-seq Studies in Crustacean Systems. Integrative and Comparative Biology 56: 1067– 1079.

DeLeo, D. M., and H. D. Bracken-Grissom, 2020 Illuminating the impact of diel vertical migration on visual gene expression in deep-sea shrimp. Mol Ecol 29: 3494–3510.

Drozdova, P., A. Kizenko, A. Saranchina, A. Gurkov, M. Firulyova et al., 2021 The diversity of opsins in Lake Baikal amphipods (Amphipoda: Gammaridae). BMC Ecology and Evolution 21: 81.

GIGA Community of Scientists, 2014 The Global Invertebrate Genomics Alliance (GIGA): Developing Community Resources to Study Diverse Invertebrate Genomes. Journal of Heredity 105: 1–18.

Gilbert, D. G., 2019 Longest protein, longest transcript or most expression, for accurate gene reconstruction of transcriptomes? bioRxiv preprint 829184.

Grabherr, M. G., B. J. Haas, M. Yassour, J. Z. Levin, D. a Thompson et al., 2011 Full-length transcriptome assembly from RNA-Seq data without a reference genome. Nature biotechnology 29: 644–52.

Guindon, S., J.-F. Dufayard, V. Lefort, M. Anisimova, W. Hordijk et al., 2010 New Algorithms and Methods to Estimate Maximum-Likelihood Phylogenies: Assessing the Performance of PhyML 3.0. Systematic Biology 59: 307–321.

Haas, B. J., A. Papanicolaou, M. Yassour, M. Grabherr, P. D. Blood et al., 2013 De novo transcript sequence reconstruction from RNA-seq using the Trinity platform for reference generation and analysis. Nature protocols 8: 1494–512.

Havird, J. C., and S. R. Santos, 2016 Here We Are, But Where Do We Go? A Systematic Review of Crustacean Transcriptomic Studies from 2014–2015. Integr Comp Biol 56: 1055–1066.

Hernández-Candia, C. N., and C. L. Tucker, 2020 Optogenetic Control of Gene Expression Using Cryptochrome 2 and a Light-Activated Degron. Methods Mol Biol 2173: 151–158.

Hitomi, K., L. DiTacchio, A. S. Arvai, J. Yamamoto, S.-T. Kim et al., 2009 Functional motifs in the (6-4) photolyase crystal structure make a comparative framework for DNA repair photolyases and clock cryptochromes. Proceedings of the National Academy of Sciences 106: 6962–6967.

Hyde, C. J., Q. P. Fitzgibbon, A. Elizur, G. G. Smith, and T. Ventura, 2020 CrustyBase: an interactive online database for crustacean transcriptomes. BMC Genomics 21: 637.

Kalyaanamoorthy, S., B. Q. Minh, T. K. F. Wong, A. von Haeseler, and L. S. Jermiin, 2017 ModelFinder: fast model selection for accurate phylogenetic estimates. Nature Methods 14: 587–589.

Kiontke, S., T. Göbel, A. Brych, and A. Batschauer, 2020 DASH-type cryptochromes – solved and open questions. Biological Chemistry 401: 1487–1493.

Kozma, M. T., H. Ngo-Vu, Y. Y. Wong, N. S. Shukla, S. D. Pawar et al., 2020 Comparison of transcriptomes from two chemosensory organs in four decapod crustaceans reveals hundreds of candidate chemoreceptor proteins. PLoS One 15: e0230266.

Le, S. Q., and O. Gascuel, 2008 An Improved General Amino Acid Replacement Matrix. Molecular Biology and Evolution 25: 1307–1320.

Leinonen, R., H. Sugawara, M. Shumway, and on behalf of the International Nucleotide Sequence Database Collaboration, 2011 The Sequence Read Archive. Nucleic Acids Research 39: D19–D21.

Lin, C., and T. Todo, 2005 The cryptochromes. Genome Biol 6: 220.

Liu, H., B. Liu, C. Zhao, M. Pepper, and C. Lin, 2011 The action mechanisms of plant cryptochromes. Trends in Plant Science 16: 684–691.

Lucas-Lledó, J. I., and M. Lynch, 2009 Evolution of Mutation Rates: Phylogenomic Analysis of the Photolyase/Cryptochrome Family. Molecular Biology and Evolution 26: 1143–1153.

Mai, U., and S. Mirarab, 2018 TreeShrink: fast and accurate detection of outlier long branches in collections of phylogenetic trees. BMC Genomics 19: 272.

Martin, J. W., and G. E. Davis, 2006 Historical Trends in Crustacean Systematics. Crustaceana 79: 1347– 1368.

Mei, Q., and V. Dvornyk, 2015 Evolutionary History of the Photolyase/Cryptochrome Superfamily in Eukaryotes. PLoS One 10: e0135940.

Minh, B. Q., M. A. T. Nguyen, and A. von Haeseler, 2013 Ultrafast Approximation for Phylogenetic Bootstrap. Molecular Biology and Evolution 30: 1188–1195.

Müller, T., and M. Vingron, 2000 Modeling Amino Acid Replacement. Journal of Computational Biology 7: 761–776.

Mykles, D. L., 2021 Signaling Pathways That Regulate the Crustacean Molting Gland. Frontiers in Endocrinology 12:.

Mykles, D. L., K. G. Burnett, D. S. Durica, B. L. Joyce, F. M. McCarthy et al., 2016 Resources and Recommendations for Using Transcriptomics to Address Grand Challenges in Comparative Biology. Integrative and Comparative Biology 56: 1183–1191.

Mykles, D. L., and E. S. Chang, 2020 Hormonal control of the crustacean molting gland: Insights from transcriptomics and proteomics. General and Comparative Endocrinology 294: 113493.

Nakasugi, K., R. Crowhurst, J. Bally, and P. Waterhouse, 2014 Combining Transcriptome Assemblies from Multiple De Novo Assemblers in the Allo-Tetraploid Plant Nicotiana benthamiana. PLoS One 9: e91776.

Nguyen, L.-T., H. A. Schmidt, A. von Haeseler, and B. Q. Minh, 2015 IQ-TREE: A Fast and Effective Stochastic Algorithm for Estimating Maximum-Likelihood Phylogenies. Molecular Biology and Evolution 32: 268–274.

Nong, W., Z. Y. H. Chai, X. Jiang, J. Qin, K. Y. Ma et al., 2020 A crustacean annotated transcriptome (CAT) database. BMC Genomics 21: 32.

Oakley, T. H., J. M. Wolfe, A. R. Lindgren, and A. K. Zaharoff, 2013 Phylotranscriptomics to Bring the Understudied into the Fold: Monophyletic Ostracoda, Fossil Placement, and Pancrustacean Phylogeny. Molecular Biology and Evolution 30: 215–233.

Oliveri, P., A. E. Fortunato, L. Petrone, T. Ishikawa-Fujiwara, Y. Kobayashi et al., 2014 The Cryptochrome/Photolyase Family in aquatic organisms. Marine Genomics 14: 23–37.

Pérez-Moreno, J. L., D. DeLeo, F. Palero, and H. D. Bracken-Grissom, 2018 Phylogenetic annotation and genomic architecture of opsin genes in Crustacea. Hydrobiologia Accepted:

Pérez-Moreno, J. L., T. M. Iliffe, and H. D. Bracken-Grissom, 2016 Life in the Underworld: Anchialine cave biology in the era of speleogenomics. International Journal of Speleology 45: 149–170.

Qin, J., Y. Hu, K. Y. Ma, X. Jiang, C. H. Ho et al., 2017 CrusTF: a comprehensive resource of transcriptomes for evolutionary and functional studies of crustacean transcription factors. BMC Genomics 18: 908.

Reitzel, A. M., L. Behrendt, and A. M. Tarrant, 2010 Light Entrained Rhythmic Gene Expression in the Sea Anemone Nematostella vectensis: The Evolution of the Animal Circadian Clock. PLOS ONE 5: e12805.

Rivera, A. S., N. Ozturk, B. Fahey, D. C. Plachetzki, B. M. Degnan et al., 2012 Blue-light-receptive cryptochrome is expressed in a sponge eye lacking neurons and opsin. Journal of Experimental Biology 215: 1278–1286.

Rozewicki, J., S. Li, K. M. Amada, D. M. Standley, and K. Katoh, 2019 MAFFT-DASH: integrated protein sequence and structural alignment. Nucleic Acids Res 47: W5–W10.

Sancar, A., 2003 Structure and Function of DNA Photolyase and Cryptochrome Blue-Light Photoreceptors. Chem. Rev. 103: 2203–2238.

Sancar, A., 2008 Structure and Function of Photolyase and in Vivo Enzymology: 50th Anniversary*. Journal of Biological Chemistry 283: 32153–32157.

Schram, F. R., 2013 Comments on Crustacean Biodiversity and Disparity of Body Plans, in Functional Morphology and Diversity, Oxford University Press, New York.

Shah, N., M. G. Nute, T. Warnow, and M. Pop, 2019 Misunderstood parameter of NCBI BLAST impacts the correctness of bioinformatics workflows (J. Hancock, Ed.). Bioinformatics 35: 1613–1614.

Simão, F. A., R. M. Waterhouse, P. Ioannidis, E. V. Kriventseva, and E. M. Zdobnov, 2015 BUSCO: assessing genome assembly and annotation completeness with single-copy orthologs. Bioinformatics 31: 3210–3212.

Smith-Unna, R., C. Boursnell, R. Patro, J. M. Hibberd, and S. Kelly, 2016 TransRate: reference-free quality assessment of de novo transcriptome assemblies. Genome Research 26: 1134–1144.

Song, L., and L. Florea, 2015 Rcorrector: efficient and accurate error correction for Illumina RNA-seq reads. GigaScience 4:.

Soubrier, J., M. Steel, M. S. Y. Lee, C. Der Sarkissian, S. Guindon et al., 2012 The Influence of Rate Heterogeneity among Sites on the Time Dependence of Molecular Rates. Molecular Biology and Evolution 29: 3345–3358.

Steenwyk, J. L., T. J. Buida, Y. Li, X.-X. Shen, and A. Rokas, 2020 ClipKIT: A multiple sequence alignment trimming software for accurate phylogenomic inference (A. Hejnol, Ed.). PLoS Biol 18: e3001007.

Stillman, J. H., J. K. Colbourne, C. E. Lee, N. H. Patel, M. R. Phillips et al., 2008 Recent advances in crustacean genomics. Integrative and Comparative Biology 48: 852–868.

Tagu, D., J. K. Colbourne, and N. Nègre, 2014 Genomic data integration for ecological and evolutionary traits in non-model organisms. BMC Genomics 15: 490.

Tang, H., R. D. Finn, and P. D. Thomas, 2019 TreeGrafter: phylogenetic tree-based annotation of proteins with Gene Ontology terms and other annotations. Bioinformatics 35: 518–520.

Tchesunov, A. V., and V. N. Ivanenko What is the difference between marine and limnetic-terrestrial associations of nematodes with invertebrates? Integrative Zoology n/a:

Teschke, M., S. Wendt, S. Kawaguchi, A. Kramer, and B. Meyer, 2011 A Circadian Clock in Antarctic Krill: An Endogenous Timing System Governs Metabolic Output Rhythms in the Euphausid Species Euphausia superba. PLOS ONE 6: e26090.

Timm, L., J. A. Browder, S. Simon, T. L. Jackson, I. C. Zink et al., 2019 A tree money grows on: the first inclusive molecular phylogeny of the economically important pink shrimp (Decapoda : Farfantepenaeus) reveals cryptic diversity. Invert. Systematics.

Waterhouse, R. M., F. Tegenfeldt, J. Li, E. M. Zdobnov, and E. V. Kriventseva, 2013 OrthoDB: a hierarchical catalog of animal, fungal and bacterial orthologs. Nucleic Acids Research 41: D358– D365.

Westerman, R., M. Ahmed, and O. Holovachov, 2022 Gammarinema scyllae sp. n. and Monhystrium mangrovi sp. n. (Nematoda: Monhysteridae) from land crabs from New Caledonia. Syst Parasitol 99: 83–101.

Wolfe, J. M., J. Luque, and H. D. Bracken-Grissom, 2021 How to become a crab: Phenotypic constraints on a recurring body plan. BioEssays 43: 2100020.

Wood, D. E., J. Lu, and B. Langmead, 2019 Improved metagenomic analysis with Kraken 2. Genome Biology 20: 257.

Wright, R. J., A. M. Comeau, and M. G. I. Langille, 2022 From defaults to databases: parameter and database choice dramatically impact the performance of metagenomic taxonomic classification tools. bioRxiv 2022.04.27.489753.

Yamada, K. D., K. Tomii, and K. Katoh, 2016 Application of the MAFFT sequence alignment program to large data—reexamination of the usefulness of chained guide trees. Bioinformatics 32: 3246– 3251.

Yang, Z., 1995 A Space-Time Process Model for the Evolution of DNA Sequences. Genetics 139: 993– 1005.

Yang, Y., Z. Cui, T. Feng, C. Bao, and Y. Xu, 2021 Transcriptome analysis elucidates key changes of pleon in the process of carcinization. J. Ocean. Limnol. 39: 1471–1484.

Zantke, J., T. Ishikawa-Fujiwara, E. Arboleda, C. Lohs, K. Schipany et al., 2013 Circadian and Circalunar Clock Interactions in a Marine Annelid. Cell Reports 5: 99–113.

Zhang, L., M. H. Hastings, E. W. Green, E. Tauber, M. Sladek et al., 2013 Dissociation of Circadian and Circatidal Timekeeping in the Marine Crustacean Eurydice pulchra. Current Biology 23: 1863– 1873.

